# Searching for the causal effects of BMI in over 300 000 individuals, using Mendelian randomization

**DOI:** 10.1101/236182

**Authors:** Louise A C Millard, Neil M Davies, Kate Tilling, Tom R Gaunt, George Davey Smith

## Abstract

Mendelian randomization (MR) has been used to estimate the causal effect of body mass index (BMI) on particular traits thought to be affected by BMI. However, BMI may also be a modifiable, causal risk factor for outcomes where there is no prior reason to suggest that a causal effect exists. We perform a MR phenome-wide association study (MR-pheWAS) to search for the causal effects of BMI in UK Biobank (n=334 968), using the PHESANT open-source phenome scan tool. Of the 20 461 tests performed, our MR-pheWAS identified 519 associations below a stringent P value threshold corresponding to a 5% estimated false discovery rate, including many previously identified causal effects. We also identified several novel effects, including protective effects of higher BMI on a set of psychosocial traits, identified initially in our preliminary MR-pheWAS and replicated in an independent subset of UK Biobank. Such associations need replicating in an independent sample.

## INTRODUCTION

Body mass index (BMI), taken to be a general indicator of adiposity, has been associated with many traits and diseases ^1–3^. These observational associations may be due to a causal effect of adiposity (as reflected in BMI), a causal effect of the phenotype on BMI, or confounding. Mendelian randomization (MR) estimates the causal effect of an exposure on an outcome using genetic variants as an instrumental variable for the exposure ^4,5^ To date, MR has been used to assess whether BMI causally affects a vast array of phenotypes ^6–21^. These results suggest that higher BMI leads to an earlier age at menarche ^9^, a reduction in physical activity ^6^, an increased risk of coronary heart Disease^22^, cancer^10,18^ and asthma^21^, a lower risk of Parkinson’s disease^17^, and changes in metabolite concentrations ^7^.

Hypothesis-free searching is an established method to identify novel associations, such as genetic variants associated with a particular phenotype in genome-wide association studies (GWAS) ^23^. In contrast to hypothesis-driven analyses, a hypothesis-free analysis can identify novel associations where there is no prior expectation that an association might exist, and should help to avoid publication bias as all results are published, not just the most “statistically significant”. Phenome scans are a class of hypothesis-free scan that test the association of a variable of interest with a potentially large array of phenotypes – the “phenome” ^24^ Phenome scans commonly seek to identify phenotypes associated with a genetic variant (phenome-wide association studies – pheWAS) or an observed phenotype (environment-wide association studies – EnWAS). A third type of phenome scan, MR-pheWAS, uses MR to search for the causal effects of a particular exposure ^25^.

To date, only a small number of phenome-wide scans of BMI have been published. We recently published a MR-PheWAS that searched for the causal effects of BMI in circa 8000 participants in the Avon Longitudinal Study of Parents and Children (ALSPAC) cohort, across 172 outcomes ^25^. This study confirmed several known associations such as with leptin levels and blood pressure, and identified potentially novel associations, such as with a self-worth score. Cronin et al. performed a pheWAS of *FTO* genetic variants within electronic health record phenotypes, in circa 25 000 participants, and identified novel associations ^26^. For instance, a genetic predisposition to a higher BMI was associated with a higher risk of fibrocystic breast disease and non-alcoholic liver disease ^26^.

UK Biobank is a prospective cohort of circa 500 000 participants ^27^, and several hypothesis-driven MR analyses of BMI have been performed in this study, to date^20,21,28,29^. The large sample size offsets the multiple testing burden of hypothesis-free analyses, providing an opportunity to search for causal effects with MR-pheWAS. Recently, we published an open-source tool for performing phenome scans (including MR-pheWAS) in UK Biobank – the PHEnome Scan ANalysis Tool (PHESANT) ^30^. PHESANT enables comprehensive phenome scans to be performed across a large and diverse set of phenotypes – all continuous, integer and categorical fields in UK Biobank – where previously researchers would restrict their analysis to a homogeneous subset of phenotypes that could be processed in a consistent fashion ^30^.

In this study we search for the causal effects of adiposity, using the PHESANT tool in UK Biobank. We use BMI as a surrogate measure of adiposity. In our presentation of PHESANT ^30^ we analyzed the initial non-random sample of circa 115 000 participants that was available in UK Biobank at that time. We found that participants with a genetic propensity to a higher BMI were less likely to perceive themselves as a nervous person or to call themselves tense or ‘highly strung’ ^30^. In this work, we search for the causal effects of BMI in circa 330 000 participants in UK Biobank satisfying our inclusion criteria. BMI is a well-studied phenotype, hence this study serves as a model for future MR-pheWAS that may investigate phenotypes with much weaker priors regarding their causal effects.

## RESULTS

We checked the strength of the association between the BMI allele score and BMI, and found a standard deviation (SD) increase in BMI allele score was associated with a 0.63 kg/m^2^ increase in BMI (95% confidence interval (CI): 0.62, 0.65, F statistic=6052.88).

### Results of MR-pheWAS analysis

The results of our MR-pheWAS include 20 461 tests ranked by P value, given in Supplementary data file 1. Figure 2 shows the number of fields reaching each stage of the PHESANT automated pipeline. A QQ plot is given in Figure 3, and the PHESANT-viz visualization can be found at [http://www.datamining.org.uk/PHESANT/results-bmi.html]. We identified 519 results at a false discovery rate of 5% (using a P value threshold of 0.05x519/20461 = 1.27×10^-3^), given in Supplementary table 4, and of these, 259 results had a P value lower than a stringent Bonferroni corrected threshold of 2.44×10^-6^ (0.05/20461).

**Figure 2:**
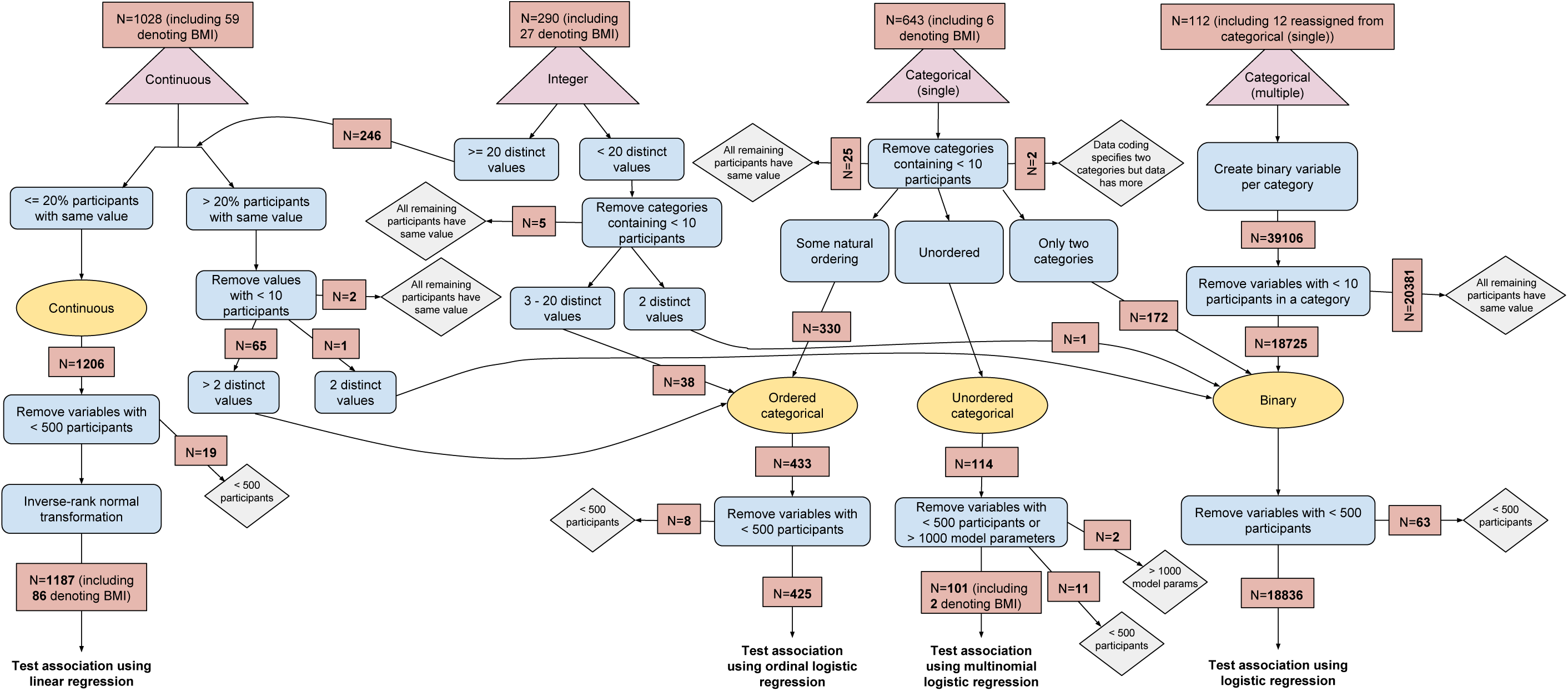
Variable processing flow diagram showing logic from defined field type in UK Biobank data to test of association, with number of variables reaching each stage of processing flow. Triangular nodes at top of figure are field types defined by UK Biobank. Rectangular nodes show processing logic used to determine the data type assignment (oval), either continuous, ordered categorical, unordered categorical or binary, and hence finally, the type of test used: linear, ordinal logistic, multinomial logistic or logistic regression, respectively. Diamond nodes show points where variables may be removed.

**Figure 3:**
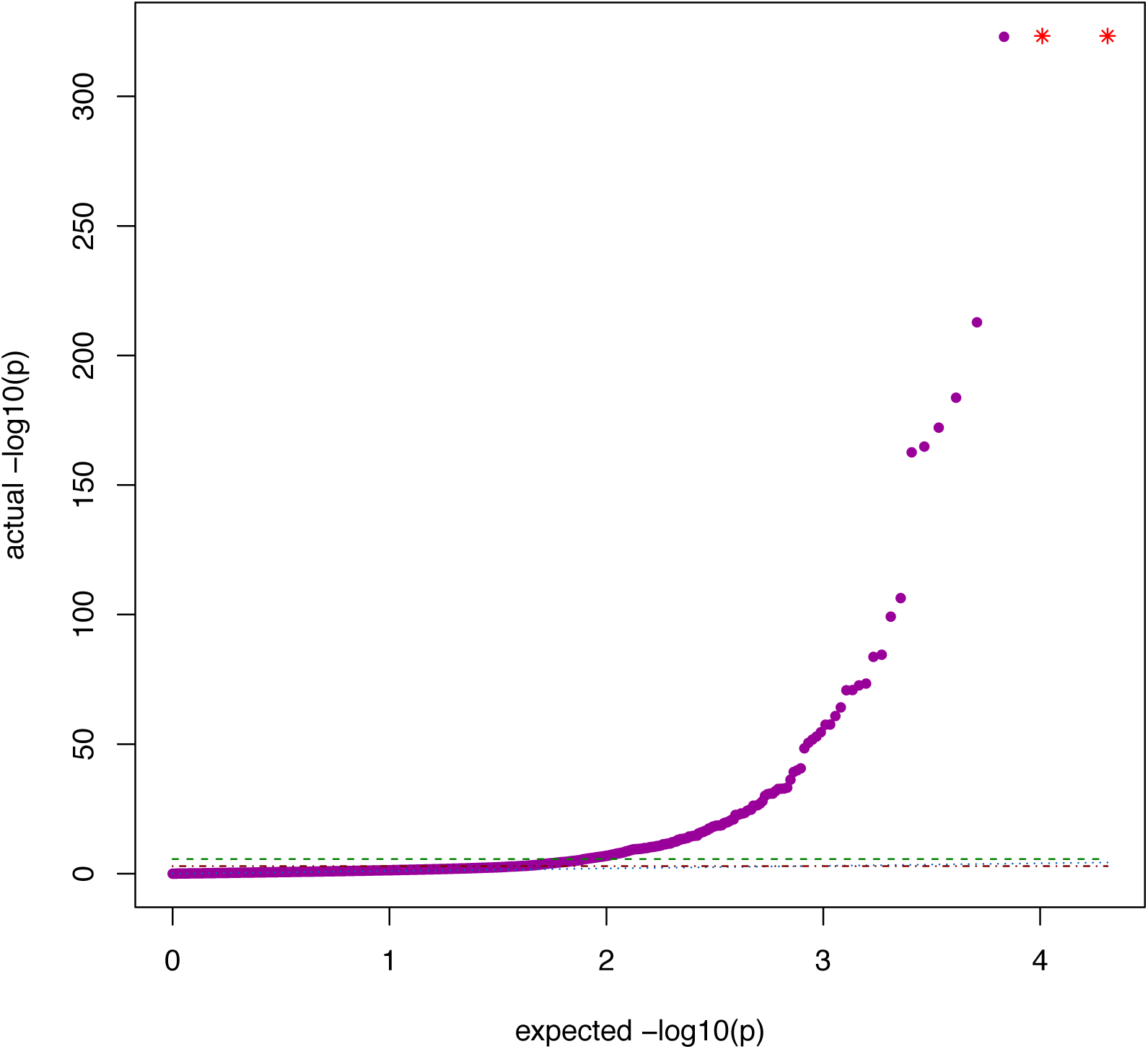
QQ plot of 20,461 MR-pheWAS results. Green dashed line: Bonferroni corrected threshold (p= 2.44x10^-6^). Red dash-dotted line: FDR threshold (p= 1.27×10^-3^). Blue dotted line: actual = expected. Purple points: results of tests performed in MR-pheWAS. Red stars: results with P values < 9.88x10^-324^.

Our MR-pheWAS identified several known effects of BMI. For example, a genetic predisposition to a higher BMI was associated with an increased risk of diabetes ^19^ (field ID (FID)=2443), hypertension ^40^ (FID=41204 value I10, and FID=4079), a higher bone mineral density ^8^ (FID=3148) and an earlier age of puberty in both sexes ^9^ (FIDs={2714, 2375, 2385}). Figure 4 shows that, when restricting to psychosocial traits, a subset is associated more strongly than would be expected by chance including several nervousness / anxiety traits.

**Figure 4:**
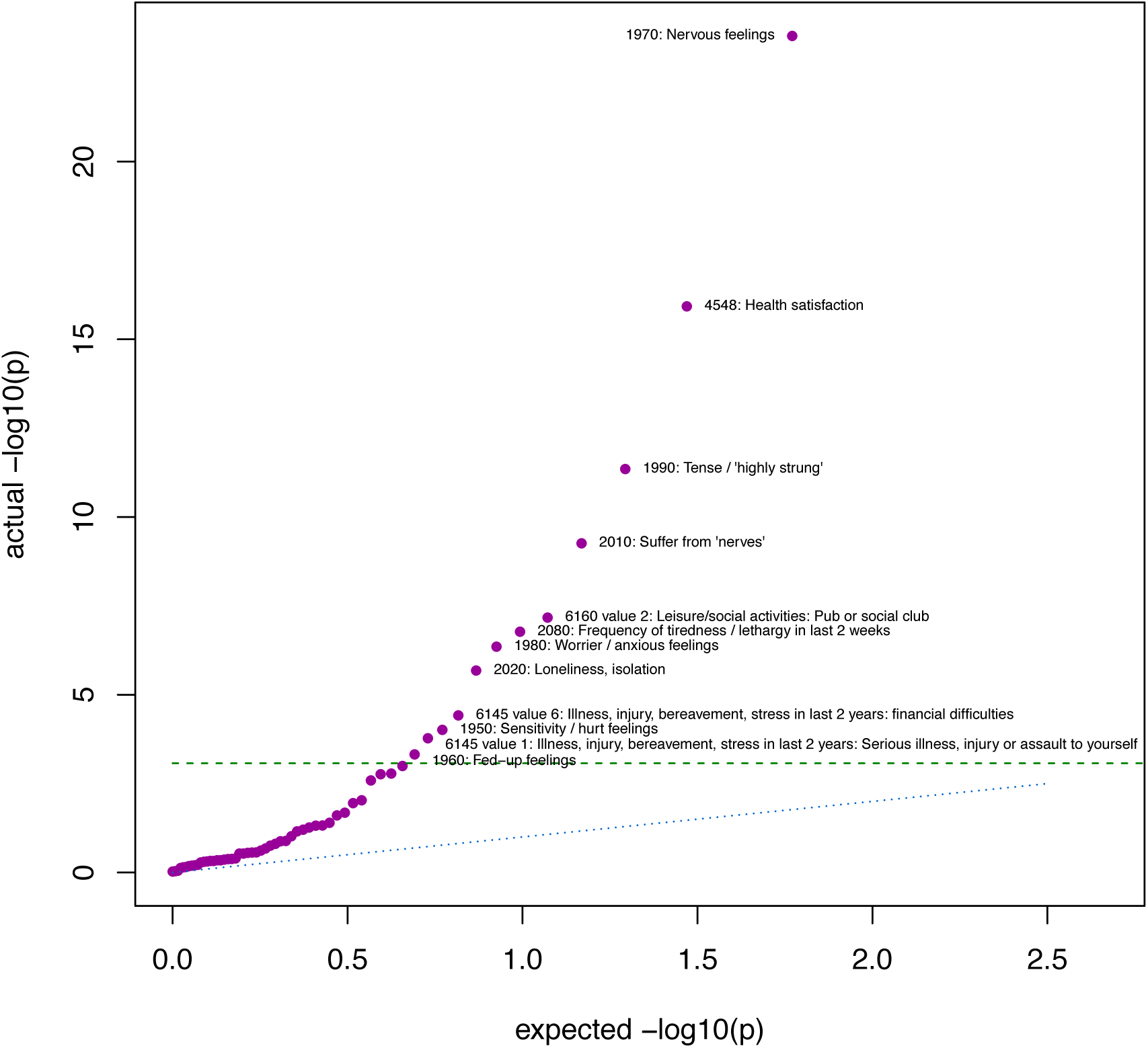
QQ plot of results in UK Biobank field category ‘psychosocial factors’ (ID 100059) Green dashed line: Bonferroni corrected threshold calculated for psychosocial traits only (0.05/59 = 8.47 × 10^-4^). Blue dotted line: actual = expected. Purple points: results of tests performed in MR-pheWAS.

#### PHESANT sensitivity analysis

Our tests of association of assessment centre with the BMI genetic score found that, while adjusting for the genetic principal components attenuated the association, an association remained (likelihood ratio test P values of 6.49×10^-26^, 4.44×10^-10^ and 6.15×10^-6^ when adjusting for age and sex only, and additionally the first 10 and first 40 genetic principal components, respectively). Supplementary figure 1 compares the results of our main MR-pheWAS analysis, adjusting for age, sex and the first 10 genetic principal components, with our sensitivity analysis, additionally adjusting for assessment centre and genetic batch. Large differences between these results often occurred for categorical outcomes with small numbers in a certain category (for multinomial logistic model this can cause a zero P value when the likelihood of the model increases to a limit ^41^; see examples in Supplementary table 5).

### Detailed follow-up of potentially novel results

Our hypothesis-free scan found that a genetic predisposition to a higher BMI was associated with a decreased reporting of being a nervous person (FID=1970, p=2.92×10^-24^), being tense or ‘highly strung’ (FID=1990, p=4.47×10^-12^), suffering from ‘nerves’ (FID=2010, p=5.47x10^-10^), or being a worrier (FID=1980, p=4.42×10^-7^). Table 1 and Supplementary figures 2 and 3 present the causal effects of BMI on these nervousness traits, estimated using two-stage instrumental variable analyses. A 1 kg/m^2^ increase in BMI caused a 6.1% [95% CI: −7.2%, −5.0%] decrease in risk of reporting as a ‘nervous’ person. Each 1 kg/m^2^ increase in BMI was observationally associated with a 3.6% [95% CI: −3.8%, −3.4%] decrease in risk of self-reporting as a ‘nervous’ person. Adjusting for age, sex and the first 40 genetic principal components did not meaningfully affect the results (see Supplementary table 6). Furthermore, causal estimates using the discovery subsample of UK Biobank (used in our initial presentation of PHESANT ^30^) and replication subsample (comprising additional participants used in this study) were consistent for these four nervousness/anxiety phenotypes.

**Table 1:** Results of follow-up analysis of nervousness / worrying traits

|  | Nervous person (FID=1970) | Being a worrier (FID=1980) | Tense / highly strung (FID=1990) | Suffering from nerves (FID=2010) |
| --- | --- | --- | --- | --- |
| N | 325,552 | 325,523 | 324,146 | 322,236 |
| **OBSERVATIONAL ESTIMATES<sup>7</sup>** |  |  |  |  |
| Observational association<sup>3*</sup> | 0.964 [0.962, 0.966] | 0.981 [0.980, 0.983] | 0.983 [0.981, 0.985] | 0.993 [0.991, 0.994] |
| **CAUSAL ESTIMATES** |  |  |  |  |
| **Estimates using two-stage IV probit regression<sup>8</sup>** |  |  |  |  |
| Effect estimate with 96 SNP score<sup>1</sup> | -0.039 [-0.047, -0.032] | -0.018 [-0.025, -0.011] | -0.028 [-0.036, -0.020] | -0.024 [-0.032, -0.017] |
| Effect estimate with 96 SNP score – approximate change of odds<sup>4*</sup> | 0.939 [0.928, 0.950] | 0.972 [0.961, 0.983] | 0.956 [0.944, 0.968] | 0.962 [0.950, 0.974] |
| Effect estimate with 96 SNP score, in terms of SD change of BMI<sup>6</sup> | -0.186 [-0.221, -0.151] | -0.085 [-0.117, -0.052] | -0.134 [-0.172, -0.096] | -0.115 [-0.152, -0.079] |
| Effect estimate with 96 SNP score, in terms of SD change of BMI, approximate change of log odds<sup>6+</sup> | -0.297 [-0.354, -0.241] | -0.135 [-0.188, -0.083] | -0.214 [-0.275, -0.154] | -0.185 [-0.243, -0.127] |
| Effect estimate with 95 SNP score (excluding FTO SNP)<sup>1</sup> | -0.040 [-0.048, -0.032] | -0.022 [-0.030, -0.015] | -0.032 [-0.040, -0.023] | -0.027 [-0.035, -0.019] |
| Effect estimate with FTO SNP<sup>1</sup> | -0.031 [-0.050, -0.012] | 0.012 [-0.006, 0.029] | -0.004 [-0.025, 0.017] | -0.008 [-0.027, 0.012] |
| **Estimates using MR-Egger<sup>2</sup>** |  |  |  |  |
| Causal effect estimate<sup>+</sup> | -0.183 [-0.405, 0.040] | -0.036 [-0.223, 0.150] | -0.123 [-0.364, 0.119] | -0.170 [-0.400, 0.060] |
| Intercept estimate | -0.002 [-0.008, 0.004] | -0.002 [-0.007, 0.003] | -0.002 [-0.009, 0.004] | 0.000 [-0.006, 0.007] |
| $I^2_{GX}$ statistic | 0.893 | | | |
| Bias adjusted causal effect estimate<sup>5+</sup> | -0.204 [-0.452, 0.043] | -0.039 [-0.248, 0.170] | -0.135 [-0.405, 0.135] | -0.193 [-0.449, 0.064] |
| **Estimates using weighted median<sup>2</sup>** |  |  |  |  |
| Weighted median effect estimate<sup>+</sup> | -0.220 [-0.315, -0.125] | -0.087 [-0.166, -0.009] | -0.172 [-0.269, -0.075] | -0.087 [-0.180, 0.007] |
| **Estimates using mode based estimate<sup>2</sup>** |  |  |  |  |
| Simple, $\phi=0.75^+$ | -0.086 [-0.317, 0.146] | -0.161 [-0.453, 0.131] | -0.273 [-0.600, 0.055] | -0.019 [-0.303, 0.266] |
| Weighted, $\phi=0.75^+$ | -0.158 [-0.286, -0.031] | 0.063 [-0.053, 0.180] | -0.114 [-0.272, 0.043] | -0.048 [-0.182, 0.085] |
| **DISCOVERY AND REPLICATION ESTIMATES<sup>9,10</sup>** |  |  |  |  |
| Discovery effect estimate with 96 SNP score<sup>1</sup> | -0.041 [-0.055, -0.027] | -0.022, [-0.035, -0.008] | -0.035 [-0.051, -0.020] | -0.029 [-0.044, -0.014] |
| Discovery Effect estimate with 96 SNP score – approximate change of odds<sup>4*</sup> | 0.936 [0.915, 0.958] | 0.966 [0.946, 0.987] | 0.945 [0.922, 0.968] | 0.955 [0.933, 0.978] |
| Replication effect estimate with 96 SNP score<sup>1</sup> | -0.040 [-0.049, -0.031] | -0.015 [-0.023, -0.006] | -0.024 [-0.034, -0.014] | -0.023 [-0.033, -0.014] |
| Replication Effect estimate with 96 SNP score – approximate change of odds<sup>4*</sup> | 0.938 [0.925, 0.952] | 0.977 [0.964, 0.990] | 0.963 [0.948, 0.978] | 0.963 [0.949, 0.978] |

BMI: body mass index; FID: field identifier; SD: standard deviation; IV: instrumental variable.
<sup>1</sup> Coefficient [95% confidence interval] from instrument variable probit models (ivprobit Stata command), for a 1 kg/m<sup>2</sup> higher BMI.
<sup>2</sup> Two sample analyses use the SNP-BMI associations from Locke GWAS<sup>33</sup>, and SNP-outcome associations estimated in UK Biobank (adjusted for and first 10 genetic principal components).
<sup>3</sup> Estimate of the odds ratio of outcome for a 1 kg/m<sup>2</sup> increase in BMI.
<sup>4</sup> Estimate of the odds ratio of outcome for a 1 kg/m<sup>2</sup> increase in BMI, calculated by taking the exponent of 1.6 times the probit estimate<sup>35</sup>.
<sup>5</sup> Bias adjusted causal effect estimate using MR-Egger with SIMEX.
<sup>6</sup> Effect estimate for SD change in BMI and logistic approximation, for comparison with two-sample approaches, as these use the BMI standardised SNP associations from the GIANT GWAS<sup>33</sup>.
<sup>7</sup> Adjusted for age and sex.
<sup>8</sup> Adjusted for age, sex and first 10 genetic principal components (see Supplementary table 6 for results of sensitivity analysis, adjusting for age, sex and first 40 genetic principal components).
<sup>9</sup> Discovery results are estimated on the UK Biobank sample used in the PHESANT application note usage example<sup>30</sup>.
<sup>10</sup> Sample size for discovery samples: Field 1970: 83,843; Field 1980: 83,795; Field 1990: 83,445; Field 2010: 82,925.
Sample size for replication samples: Field 1970: 216,311; Field 1980: 216,301; Field 1990: 215,435; Field 2010: 214,152.
<sup>+</sup> Estimates are in terms of change of log odds of outcome variable for a 1 SD higher BMI.
<sup>\*</sup> Estimates are in terms of odds ratio of outcome variable for a 1 kg/m<sup>2</sup> higher BMI.

As a sensitivity analysis, we estimated causal effects of BMI using two-sample MR methods. Supplementary figures 4 and 5 show the MR-Egger regression and simulation extrapolation (SIMEX) plots, respectively. We found little evidence of directional pleiotropy, as all confidence intervals for intercept estimates of MR-Egger included zero. While there was evidence that the ‘NO Measurement Error’ (NOME) assumption was not fully satisfied (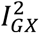 statistic = 0.893), estimates using MR-Egger with and without SIMEX were highly consistent.

Supplementary figures 6 and 7 show the smoothed empirical densities of the SNP effect estimates using mode-based estimator (MBE) smoothing parameter ϕ values of 0.5, 0.75 and 1. We choose to use ϕ=0.75 as this provided sufficient smoothing while still allowing multiple peaks to be captured. The effect estimates across IV methods – IV probit regression, and the two-sample approaches; MR-Egger, weighted median and MBE – were broadly consistent. We did, however, see a difference between the estimates of the IV probit regression and weighted MBE for reporting ‘being a worrier’ (FID=1980), where a 1 SD higher BMI was associated with a 12.7% [95% CI: −17.1%, −8.0%] decrease in the risk of reporting ‘being a worrier’ using IV probit regression, but a 6.5% [95% CI: −5.2%, 19.7%] increase in risk of reporting ‘being a worrier’ using the weighted MBE (*ϕ*=*0.75*). Furthermore, the causal estimates using the 95 SNP score versus a BMI-associated SNP in *FTO* (*“FTO* SNP”) were broadly consistent for the outcomes in our follow-up analysis, except for reporting ‘being a worrier’ (FID=1980).

The results of these four nervousness/anxiety traits do not provide a form of replication for each other because these traits are highly correlated (see Supplementary table 7; all accuracy between 57.59% and 83.49%) and so are more likely to agree compared to tests using independent traits or a replication on an independent dataset.

We searched for other nervousness related traits by searching for the terms: “worr”, “nerv”, “tens”, “anxi”, in our results listing, to determine if other related traits were tested and an association not identified, to assess the strength of the evidence when considering results for all similar phenotypes. This identified three self-reported phenotypes describing: 1) whether the participant has seen a psychiatrist for nerves, anxiety, tension or depression, (FID=2100; estimated odds ratio of 1.009 [95% CI: 0.994, 1.023]), 2) whether the participant has seen doctor (GP) for nerves, anxiety, tension or depression” (FID=2090; estimated odds ratio of 1.006 [95% CI: 0.994, 1. 017]), and 3) the frequency of tenseness / restlessness in last 2 weeks (FID=2070; not at all versus several days or more; estimated odds ratio of 0.984 [95% CI: 0.973, 0.996]).

## DISCUSSION

In this study we used the PHESANT phenome scan tool to search for the causal effects of BMI – a MR-pheWAS analysis – in a sample of circa 330 000 participants in UK Biobank, across over twenty thousand diverse phenotypes. This systematic approach helps to avoid biases associated with hypothesis-driven analyses, where a researcher might try several tests of association until a desired result is found, and should help to avoid publication bias as all results are published, not just the most “statistically significant” ^42^.

Our results include associations consistent with previous MR studies, such as adverse effects of higher BMI on risk of diabetes ^19^ and hypertension ^40^, a higher bone mineral density ^8^ and an earlier age of puberty in both sexes ^9^. Consistent with our preliminary results ^30^, we identified an association with a set of nervousness phenotypes, where a genetic predisposition to a higher BMI was associated with a person being less likely to call themselves a nervous person, tense or highly strung, a worrier, or to report they suffer from ‘nerves’. We followed up this analysis to estimate the causal effect and found that each 1 kg/m^2^ decrease in BMI caused a 6.1% [95% CI: 5.0%, 7.2%] increase in likelihood of self-reporting being a nervous person. The causal estimates were more extreme than the observational estimates for three of these outcomes. Furthermore, causal estimates using the discovery and replication subsamples of UK Biobank were consistent for these four nervousness/anxiety phenotypes.

We searched our results for other nervousness traits to determine the strength of the evidence in the context of the results of related phenotypes, which, while not a form of replication, provides an unbiased view of the evidence using the UK Biobank cohort. We identified two phenotypes detailing treatment by a psychiatrist or a doctor, respectively, for nerves, anxiety, tension or depression, but we did not find evidence of a causal effect of BMI on these phenotypes. A third result, the frequency of tenseness / restlessness in the last 2 weeks, was weakly associated in a direction consistent with the other nervousness associations we identified, although it did not pass the P value threshold corresponding to an estimated FDR of 5%.

Several previous observational studies have reported the association of anxiety and BMI. The prevalence of anxiety has been shown to be higher in obese compared with non-obese people ^43–47^, although Vainik et al. found no correlation between BMI and neuroticism ^48^. The few instrumental variable studies estimating the causal effect of BMI on anxiety that have been performed to date (one specifically looking at phobic anxiety ^49^ and the other using an anxiety measure defined using the Hospital Anxiety and Depression Scale (HADS) ^50^), did not find evidence of a causal effect, although this may be due to insufficient statistical power ^49,50^. Two recent MR studies provided evidence that an increase in BMI adversely affects risk of depression symptoms ^51^ and major depressive disorder ^52^. Furthermore, it has been shown that the mechanism through which BMI-associated genetic variants affect risk of obesity may involve regions of the brain such as the basal ganglia ^53^, which are plausibly involved in emotional processes ^54^.

The associations we identified with self-reported nervousness should be further investigated and replicated in an independent sample. These associations may reflect a true causal effect of BMI. Alternative explanations for these results include chance, or because some variants in the BMI genetic score have horizontal pleiotropic effects and are thus invalid instruments for BMI. We assessed the possibility that our instrument is invalid by comparing the estimates of two independent genetic instruments (i.e. *FTO* SNP versus the remaining genetic variants), and using alternative approaches that estimate the causal effects under differing assumptions of instrument validity – MR-Egger, weighted median and MBE. While we found little evidence of directional pleiotropy, we did find evidence of non-directional pleiotropy for the ‘being a worrier’ outcome, indicated by different estimates using the 95 SNP score versus *FTO* SNP, and also different estimates using IV probit regression compared to the weighted MBE.

UK Biobank is a highly-selected sample of the UK population, having a response rate of 5.5% ^55^, that is not representative of the UK general population ^56^. For example, UK Biobank participants have, on average, a lower BMI, and fewer self-reported health conditions, compared with the general population ^56^. Hence, our estimates may be biased if selection into the sample is affected by BMI ^57^ Also, if selection is additionally dependent on a given outcome (e.g. self-reported nervousness), associations may be biased by a particular form of collider bias – selection induced collider bias ^58^. In general, collider bias may occur when two variables (A and B) independently affect a third variable (C) and variable C is conditioned upon in analyses. Selection induced collider bias may occur when variable C represents whether a person is selected into the sample, i.e. variables A and B both independently affect participation in the study. Hence, estimates of association between two phenotypes – such as our BMI genetic score and a given outcome in our study – can be biased, if inclusion in the study is affected by both phenotypes.

We found that our BMI genetic score was associated with assessment centre, even after adjustment for the first 40 principal components. This may indicate that the genetic principal components are not fully accounting for genetic population differences. However, this may also be due to selection induced collider bias, because both BMI and location are related to selection into the sample ^56^. If both BMI and assessment centre affect participation in the study, then an association may be induced between the BMI genetic score and assessment centre. For example, the South West region had the highest participation rate of the regions sampled, such that living in this region is associated with a higher chance of participating compared to the other regions ^56^. Since BMI is negatively associated with participation in UK Biobank, we would expect (under most realistic assumptions about the association between BMI, location and participation in UK Biobank) to see a positive association between the BMI genetic risk score, and participating in the South West region compared to other regions if collider bias is the cause, and (using attendance at the Bristol assessment centre as a proxy for region of residence) this is indeed what we see (see illustration shown in Supplementary figure 8). Furthermore, the West Scotland region has the lowest participation rate, and attending the Glasgow assessment centre compared to other centres is associated with a lower BMI genetic risk score, which is expected if this relationship is due to selection induced collider bias.

We now discuss some further limitations of this work. PHESANT uses a rule-based method to automatically determine how to test each outcome, and it is possible that this may deal with some variables inappropriately. Also, PHESANT tests the linear association of the genetic score with the set of outcomes, and it is possible that nonlinear associations may exist. For instance, it is possible that BMI has a non-linear effect on nervousness/anxiety (for example low and high BMI may cause higher levels of anxiety compared with those in the normal range) and in follow-up analyses this should be investigated. Ranking the associations means that we should expect the true strength of the associations to be less than we reported due to the winner’s curse. We used stringent P value thresholds, which, although reducing the type I error rate, is likely to also increase the type II error rate. We used BMI as a surrogate measure for adiposity, but the effect of adiposity estimated using observational data is not the true effect of an intervention that modifies BMI, because this will depend on the particular intervention used. The causal effects we have estimated may be the result of changes of other aspects of BMI in addition to or instead of changes in adiposity (e.g. changes in lean mass versus fat mass) ^59^. Similarly, our BMI genetic score is an instrument for life-long BMI, hence we cannot say that it is BMI at a specific age (e.g. the age of the UK Biobank cohort) that affects an outcome. However, searching for potential causal effects using MR is useful to identify outcomes that may be modified through interventions on modifiable determinants of adiposity (e.g. diet or physical activity) ^59^.

PHESANT is the first tool to perform comprehensive phenome scans, where previously the set of outcomes tested would be restricted to a homogeneous subset ^30^. In this study, we have presented the first comprehensive phenome scan, using the PHESANT tool to search for the causal effects of BMI. This MR-pheWAS confirmed several established effects of BMI, and also identified potentially interesting novel results, such as a potential causal effect of BMI on feelings of nervousness. This work demonstrates how MR-pheWAS can be applied in UK Biobank and can serve as a model for future studies. There is much potential to use MR-pheWAS to search for the causal effects of phenotypes where, compared to BMI, much weaker priors regarding their causal effects. Phenome-wide scans are a hypothesis generation approach and hence identified associations should be followed-up in an independent sample.

## METHODS

### Study population

UK Biobank sampled 503 325 men and women in the UK aged between 37–73 years (99.5% were between 40 and 69 years) ^31^. This cohort includes a large and diverse range of data from blood, urine and saliva samples and health and lifestyle questionnaires ^27^

Of the 487 406 participants with genetic data, we removed 373 with genetic sex different to reported sex, and 471 with sex chromosome aneuploidy (identified as putatively carrying sex chromosome configurations that are not either XX or XY). We found no outliers in heterozygosity and missing rates, which would indicate poor quality of the genotypes. We removed 78 309 participants not of white British ancestry^32^. We removed 73 277 participants who were identified as being related, having a kinship coefficient denoting a third degree (or closer) relatedness ^32^. We removed 8 individuals with withdrawn consent, giving a sample of 334 968 participants. A participant flow diagram is given in Figure 1.

**Figure 1:**
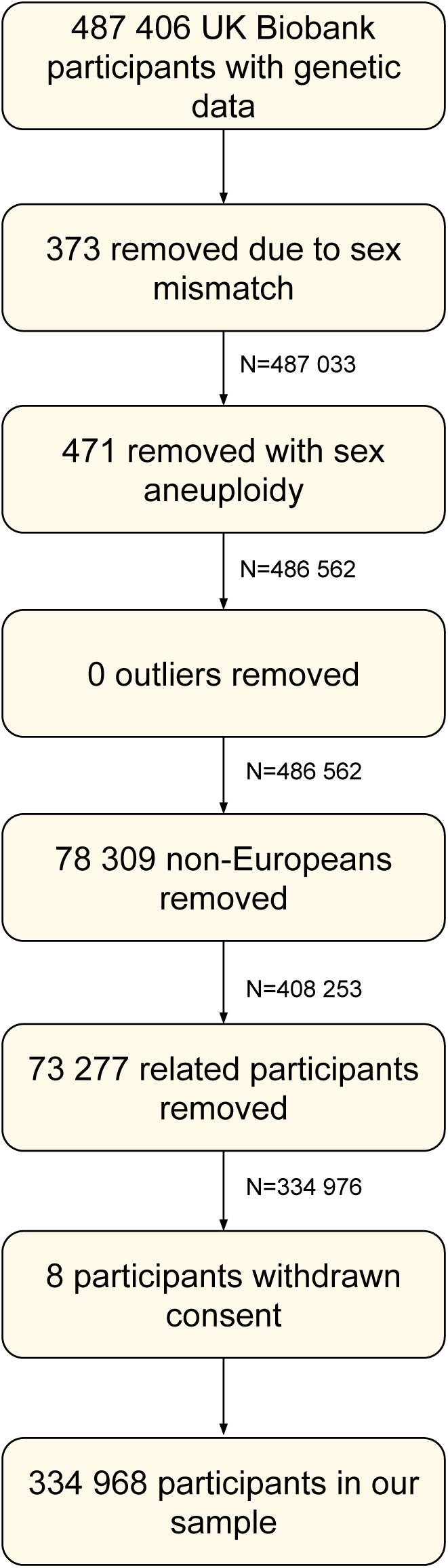
Participant flow diagram.

### BMI allele score

We created an allele score from 96 genetic variants previously found to be associated with BMI, in a recent GWAS meta-analysis by the GIANT consortium ^33^. The score was calculated as a sum of the number of BMI-increasing alleles, weighted by the effect size as reported in the GIANT GWAS (reported as a SD change of BMI per dosage increase) ^33^ (see Supplementary table 1), such that a higher allele score corresponds to a higher BMI, and was standardized to have a mean of zero and SD of 1.

### Outcomes

The Biobank data showcase allows researchers to identify variables based on the field type (http://biobank.ctsu.ox.ac.uk/showcase/list.cgi). At the time of initial data download there were 2143 fields of the following types: integer, continuous, categorical (single) and categorical (multiple) (identified in March 2016, and updated in November 2017 for fields where data accrual was still underway).

We excluded 70 fields a priori, given in Supplementary table 2, for the following reasons. We removed one field denoting the assessment centre. We removed 2 fields described by UK Biobank as ‘polymorphic’, containing values with mixed data types. We removed 14 fields that, although listed in the data showcase, were not currently available. We removed 13 genetic descriptor fields, 1 sex field and 4 age fields. We removed 17 variables describing the assessment centre environment. We removed 18 categorical (single) fields with more than one value recorded per person.

We assigned 92 outcomes that denote adiposity or some aspect of weight, fat mass or height as ‘exposure phenotypes’ – to be included in our MR-pheWAS analysis so that we can assess the strength of outcome associations in relation to these, while this a priori assignment means they will not contribute to the multiple testing burden (see Supplementary table 3).

This resulted in a set of 1981 UK Biobank fields (263 integer, 969 continuous, 649 categorical (single) and 100 categorical (multiple)), referred to hereafter as the outcome dataset (because they are tested as an outcome irrespective of whether this is biologically plausible).

### Observed BMI

Weight and height were measured at the initial UK Biobank assessment centre – weight in light clothing and unshod was measured using Tanita BC418MA body composition analyser to the nearest 100g, and height to the nearest cm using at Seca 202 device. These were used to calculate BMI (kg/m^2^).

### Covariates

We include age and sex as covariates in our models to reduce the variation in our outcomes. Age when participants attended the UK Biobank assessment centre was derived from their date of birth and the date of their assessment centre visit. Sex was self-reported during the touchscreen questionnaire (and validated using the genome-wide data). We adjust for the first 10 genetic principal components to control for confounding via population stratification. Genetic variants are set at conception, and after conception they cannot be affected by traditional confounding factors, therefore we did not adjust for any further covariates.

### Statistical methods

#### PHESANT MR-pheWAS

We test the direct association of the BMI genetic score with each of the outcome variables using the PHESANT package (version 0.10). A description of PHESANT’s automated rule-based method is given in detail elsewhere ^30^. In brief, the decision rules start with the variable field type and use rules to categorize each variable as one of four data types: continuous, ordered categorical, unordered categorical or binary. Variables with the continuous and integer field type are usually assigned to the continuous data type, but some are assigned to ordered categorical if, for instance, there are only a few distinct values. Variables of the categorical (single) field type are assigned to either the binary, ordered categorical or unordered categorical, depending on whether the field has two distinct values, or has been specified as ordered or unordered in the PHESANT setup files. Variables of the categorical (multiple) field type are converted to a set of binary variables, one for each value in the categorical (multiple) fields.

PHESANT estimates the bivariate association of the BMI genetic score with each outcome variable. The BMI genetic score and outcome variables are the independent (exposure) and dependent (outcome) variables in the regression model, respectively. Outcome variables with continuous, binary, ordered categorical and unordered categorical data types, are tested using linear, logistic, ordered logistic, and multinomial logistic regression, respectively. Prior to testing, an inverse normal rank transform is applied to variables of the continuous data type, to ensure they are normally distributed. All analyses are adjusted for covariates as described above.

We correct for multiple testing by controlling for the expected proportion of false positive results. After ranking the results by P value, we identify the largest rank position with a P value less than *P_threshoid_* = 0.05×rank/n, where *n* is the total number of tests in the phenome scan. *P_threshoid_* is the P value threshold resulting in a false discovery rate of 5% ^34^ We also calculate a stringent Bonferroni corrected P-value threshold, by dividing 0.05 by the number of tests performed, which assumes each test is independent.

##### Results visualization with PHESANT-viz

We use PHESANT-viz, a D3 Javascript visualization tool included in the PHESANT package, to visualize our results as a graph, using the Biobank assigned category structure (http://biobank.ctsu.ox.ac.uk/showcase/label.cgi). This enables interpretation of the identified associations with consideration of the results for related variables. For example, the estimated effects of BMI on depression are grouped with the results of other psychosocial phenotypes.

##### PHESANT sensitivity analysis

We re-run our phenome scan to assess residual confounding, additionally adjusting for both assessment centre and genetic batch.

#### Testing for residual population stratification

We test the extent that genetic principal components account for genetic differences across the population. We test the association of assessment centre with the BMI genetic score (as the independent and dependent variables, respectively), adjusting for:

1. Age and sex.
2. Age, sex and the first 10 genetic principal components.
3. Age, sex and the first 40 genetic principal components.

We use a likelihood ratio test to determine the strength of the association of the assessment centres collectively, with the genetic score. An association between the assessment centres and the BMI genetic score, after adjusting for the genetic principal components, may indicate that the genetic principal components are not fully accounting for population stratification.

#### Follow-up analysis of identified associations

We identified associations with a related set of psychosocial traits and generate a QQ-plot restricting to the psychosocial UK Biobank category only (category ID=100059), to determine whether these results have an association stronger than expected by chance, given the results of related phenotypes. We perform a formal instrumental variable analysis for a set of binary outcomes, using two-stage IV probit regression (the Stata `ivprobit` command, with conditional maximum-likelihood estimation), adjusting for age, sex and the first 10 genetic principal components. We also adjust for age, sex and the first 40 genetic principal components, as a sensitivity analysis. We take the exponent of 1.6 times the estimates, to approximate the association in terms of the change of odds ^35^. We also test the observational association of BMI with each phenotype using logistic regression (Stata `logistic` command).

We compare estimates within the initial discovery sample (used in our initial presentation of PHESANT ^30^), with estimates using the additional participants used in this study, as a replication. We remove all participants from our replication sample who were identified as being related to a participant in the discovery sample, having a kinship coefficient denoting a third degree (or closer) relatedness.

##### Follow-up sensitivity analyses

We performed sensitivity analyses to investigate whether our genetic score, and its constituent genetic variants, may be invalid instruments for BMI. First, we compared the effect estimates using two independent instrumental variables: 1) SNP rs1558902 (the SNP most strongly associated with BMI, at the *FTO* locus), and 2) the remaining 95 genetic variants.

Second, we also estimated causal effects using three alternative MR approaches – MR-Egger^36^, weighted median^37^ and MBE^38^. These methods estimate effects consistent with the true causal effect under more relaxed assumptions of instrument validity, compared with IV probit regression where for example, including just one genetic variant with pleiotropic effects could bias estimates. These methods are performed using two sample MR, using estimates of association of each genetic variant on BMI, and estimates of association of each genetic variant on a given outcome, respectively, estimated in different populations. We use the SNP-BMI estimates from the GIANT GWAS ^33^ (that did not include UK Biobank), and estimate the SNP-outcome associations on our UK Biobank sample, adjusting for age, sex and the first 10 genetic principal components.

The MR-Egger, weighted median and MBE approaches are complementary, each depending on distinct assumptions about instrument validity. Estimates from MR-Egger are not biased by horizontal pleiotropy, where a genetic variant affects several traits through separate pathways, under the assumption that the association of each genetic variant with the exposure is independent of any horizontal pleiotropic effect – referred to as the InSIDE (Instrument Strength Independent of Direct Effects) assumption ^36^. This approach can test for directional pleiotropy, which is when the horizontal pleiotropic effects of the genetic variants are not balanced about the null ^36^. Directional pleiotropy is identified where the intercept estimate is not consistent with the null. MR-Egger assumes that the SNP-exposure associations are measured without error, known as the NOME assumption ^39^. We determine the degree to which the NOME assumption is violated, using the 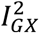 statistic, which is an adaption of the *I^2^* statistic used in the field of meta-analysis ^39^. The 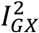 statistic ranges between 0 and 1, and captures the uncertainty in the estimated SNP-exposure associations relative to the heterogeneity across the true underlying SNP-exposure associations. It is recommended that results of MR-Egger should be treated with caution when 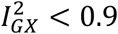 ^36^. Furthermore, we performed an MR-Egger analysis with bias adjustment using SIMEX, to estimate the causal effect when the NOME assumption is violated ^39^. In brief, SIMEX works by learning models with increasing violations of the NOME assumption (i.e. increasing error in the SNP-exposure associations). The estimates from these models are then treated as a set of data points and a model is learnt across these. This ‘meta’ model is then used to extrapolate back to the estimate that would have occurred if the NOME assumption was satisfied.

The weighted median approach estimates a consistent causal effect under the assumption that less than 50% of the genetic variants are invalid instruments ^37^ The simple version of MBE assumes that the mode of the smoothed empirical density function of SNP estimates is consistent with the true causal effect, even if the majority of the SNP estimates are not consistent – known as the ZEro Mode Pleiotropy Assumption (ZEMPA) ^38^. We also test using a weighted version of the MBE, which instead assumes that the mode of the inverse-variance weighted empirical density function is consistent with the true causal effect. The MBE method uses a bandwidth parameter *φ,* which determines the amount of smoothing of the empirical density. We generate the smoothed empirical density using a range of *ϕ* values, and choose a *ϕ* value that provides an appropriate amount of smoothing (a degree of smoothing that allows multi-modal distributions to be identified while not overfitting to the estimated values). We test the simple and weighted versions of the MBE using our chosen *ϕ* value.

Analyses are performed in R version 3.2.4 ATLAS, Matlab r2015a or Stata version 14, and code is available at [https://github.com/MRCIEU/PHESANT-MR-pheWAS-BMI]. Git tag v0.2 corresponds to the version presented here.

## ACKNOWLEDGEMENTS

This work was supported by the University of Bristol and UK Medical Research Council [grant numbers MC_UU_12013/1, MC_UU_12013/8 and MC_UU_12013/9]. LACM is funded by a University of Bristol Vice-Chancellor’s Fellowship. This research has been conducted using the UK Biobank Resource under Application Number 16729.

## AUTHOR CONTRIBUTIONS

LACM contributed to the design of the study, performed all analyses, wrote the first version of the manuscript, critically reviewed and revised the manuscript and approved the final version of the manuscript as submitted. NMD contributed to the design of the study, critically reviewed and revised the manuscript and approved the final version of the manuscript as submitted. KT contributed to the design of the study, critically reviewed and revised the manuscript and approved the final version of the manuscript as submitted. TRG critically reviewed and revised the manuscript and approved the final version of the manuscript as submitted. GDS conceptualized the study, contributed to the design of the study, critically reviewed and revised the manuscript and approved the final version of the manuscript as submitted.

